# The biogeography of hydrothermal vent giant viruses across the global ocean

**DOI:** 10.64898/2026.05.12.724634

**Authors:** Md Moinuddin Sheam, Elaine Luo

## Abstract

Viruses are the most abundant known biological entities in the ocean and play central role in microbial ecology. However, the patterns and drivers of their biogeographic distribution across global ocean remain poorly understood. Here, we conducted large-scale metagenomic analysis of 514 samples from globally-distributed hydrothermal vents and the pelagic ocean to examine the biogeography of giant viruses, a widespread viral group in the ocean. We identified 279 putative giant virus populations from hydrothermal vents and 56 populations were broadly distributed across hydrothermal vents. Vent-derived giant viruses are also present in the global pelagic ocean showing an “everything is everywhere” biogeographic pattern. We identified the mechanistic processes governing this global distribution of giant viruses and found that horizontal passive transport, mediated by hydrothermal plumes, is likely a major mechanism that drives the global connectivity of vent-derived giant viruses. Additionally, environmental selection appears to shape viral distributions within high-temperature vent habitats, as giant viruses encoding heat-shock proteins were enriched in these environments. Overall, this study identified the key process that sustain global-scale ecological connectivity for viruses between hydrothermal vents and pelagic ocean and demonstrates that a classic microbial biogeography paradigm, “everything is everywhere, but the environment selects” applies to marine giant viruses.

## Introduction

The Baas Becking hypothesis, which states “everything is everywhere, but the environment selects”, frames our understanding of the spatial distribution of microbes across our globe. The “everything is everywhere” aspect highlights the remarkable dispersal potential of microorganisms [1]. In hydrothermal vent systems, geographically-isolated hotspots of primary production near the seafloor, bacterial genera considered to be endemic were also found more broadly dispersed in the open ocean water column [2] and certain archaeal strains demonstrated widespread dispersal capacity across geographically-distant vents [3]. Hydrothermal plumes can disperse horizontally for hundreds of kilometers through deep-ocean currents [4–6]. Such plume-mediated dispersal can facilitate the long-distance spatial distribution of vent-derived microorganisms [2], as well as planktonic larvae [7], contributing to ecological connectivity between pelagic and vent habitats. This potential for environmental connectivity (“everything is everywhere”) and unique geochemical characteristic (“environment selects”) makes hydrothermal vents a highly suitable system for elucidating ecological connectivity between vent and pelagic environments, as well as the mechanistic processes governing microbial distribution in the marine ecosystem.

Whether the Becking hypothesis can also be applied to marine viruses, the most abundant known biological entities on our planet with an estimated 10^30^ viruses in the global ocean [8], remains open to investigation. Viruses are abundant in hydrothermal vent environments, and virus-like particles range in abundance from 10 to 10 /mL in samples collected from actively vent sites [9]. Previous work on viruses inhabiting hydrothermal vents, focused primarily on bacteriophages, found that viral populations were largely endemic to the vent site and habitat type [10]. For instance, a metagenomic survey of viral communities from hydrothermal vents across Axial Seamount (Pacific Ocean) and Mid Cayman Rise (Atlantic) reported largely endemic viral populations, indicating limited dispersal amongst vent sites [11]. A similar pattern of endemic distribution was observed for viruses across globally distributed hydrothermal vents [12]. Whether this observed endemism of vent-derived bacteriophage populations can be applied more broadly to giant viruses (phylum *Nucleocytoviricota)* remains open to investigation, as aside from a few studies [13–15], giant viruses remain largely understudied in hydrothermal vent ecosystems.

Giant viruses, or nucleocytoplasmic large DNA viruses (NCLDVs), are diverse and abundant in the marine environment, with an estimated 10^4^–10^6^ giant viruses per milliliter of seawater [16]. Their size can reach up to 1.5 μm, and their genomes can reach up to ∼2.5 million base pairs [17]. We postulate that, relative to bacteriophages, the large genome sizes and the resulting functional versatility of giant viruses may facilitate a wider distribution and adaptation potential across the ocean. Giant viruses are distributed across different environments, ranging from marine, freshwater, terrestrial, wastewater, thermal spring, and hydrothermal vents [15]. While metagenomics and culture-based studies have identified the presence of giant viruses in hydrothermal vents [14,15], the biogeography of vent-derived giant viruses and how it relates to the Becking hypothesis remains open to investigation. Here, we aim to identify the patterns and drivers of the biogeography of giant viruses to identify their dispersal capacity and the role of environmental selection in shaping the distribution of this key group of microorganisms across the global ocean.

In this study, we conducted a large-scale metagenomic analysis of 514 samples across six distinct, globally-distributed hydrothermal vents to identify the patterns and drivers of the biogeography of giant viruses. We observed a wide distribution of giant viruses across globally-distributed vent sites, with phylogenetically-similar giant viruses detected across geographically-distant hydrothermal vents. This broad distribution pattern contrasts with the site-specific endemism previously observed in bacteriophages. To further identify the mechanism behind the widespread distribution of giant viruses, we looked for evidence of the distribution of vent–derived giant viruses beyond vent ecosystems by analyzing an additional 868 metagenomic samples obtained from the pelagic global ocean. We found that giant viruses originating from hydrothermal vents were widely distributed in the global ocean, likely reflecting horizontal passive transport through hydrothermal plumes. We show that, for giant viruses, “everything is everywhere,” driven by their high dispersal potential through plume-mediated transport.

Additionally, we observed evidence supporting local thermal adaptation and that “the environment selects” for vent-adapted giant viruses in high-temperature habitat. Overall, our study reveals the cosmopolitan distribution of hydrothermal vent giant viruses across the global ocean, likely facilitated by their metabolic flexibility and plume-mediated transport, aligning with the Bass Becking hypothesis of microbial biogeography. This work opens new paradigms in viral ecology, revealing global-scale ecological connectivity between deep-sea hydrothermal vents and the pelagic ocean.

## Results and discussions

### Presence of giant viruses in deep-sea hydrothermal vents

We curated a database of 279 nonredundant NCLDV contigs (>5 kbp) based on NCLDV-specific marker genes from deep-sea hydrothermal vent metagenomes. Across six vent locations, approximately 0.03–1.5% of reads mapped to these contigs, with >1% mapping at Axial Seamount, Lau Basin, Mid Cayman Rise, and Southern Mariana (fig. S2). Diffuse fluid samples showed a higher proportion of reads (∼1.5%) mapping to NCLDV contigs compared to plume and hydrothermal fluid samples (fig. S3), likely reflecting the presence of diverse microbial communities in diffuse fluids driven by mixing of hydrothermal fluids with seawater [18]. Most of the vent giant viruses were taxonomically novel. Order-level taxonomy was assigned to 27% of contigs, revealing *Imitervirales*, *Pimascovirales*, and *Algavirales*, with members of *Imitervirales* being particularly prominent (fig. S4, table S2) .Members of these orders primarily infect heterotrophic protists and algae [16], suggesting that the giant viruses identified in hydrothermal vent environments may infect microbial eukaryotic hosts.

### Wide biogeographical distribution of giant viruses across hydrothermal vents

Giant viruses exhibited broad biogeographical distribution across six geographically distinct vent locations (Fig. 1). Among the 279 vent-derived NCLDV populations, 56 were detected across at least four vent systems, and three contigs were detected in all six (Fig. 2a, table S3). All widely-distributed populations (defined as present across four or more locations) were taxonomically novel (fig. S5). Samples from the Axial Seamount and Lau Basin, both located in the Pacific Ocean, shared the highest number of giant virus contigs. In contrast, 31 contigs were detected exclusively in the Axial Seamount, indicating the presence of endemic viral populations unique to this vent system. The biogeographic patterns of giant viruses in hydrothermal vents should be interpreted qualitatively rather than quantitatively, as variation in the number of metagenomic samples across vent systems can affect contig recovery.

**Fig. 1.**
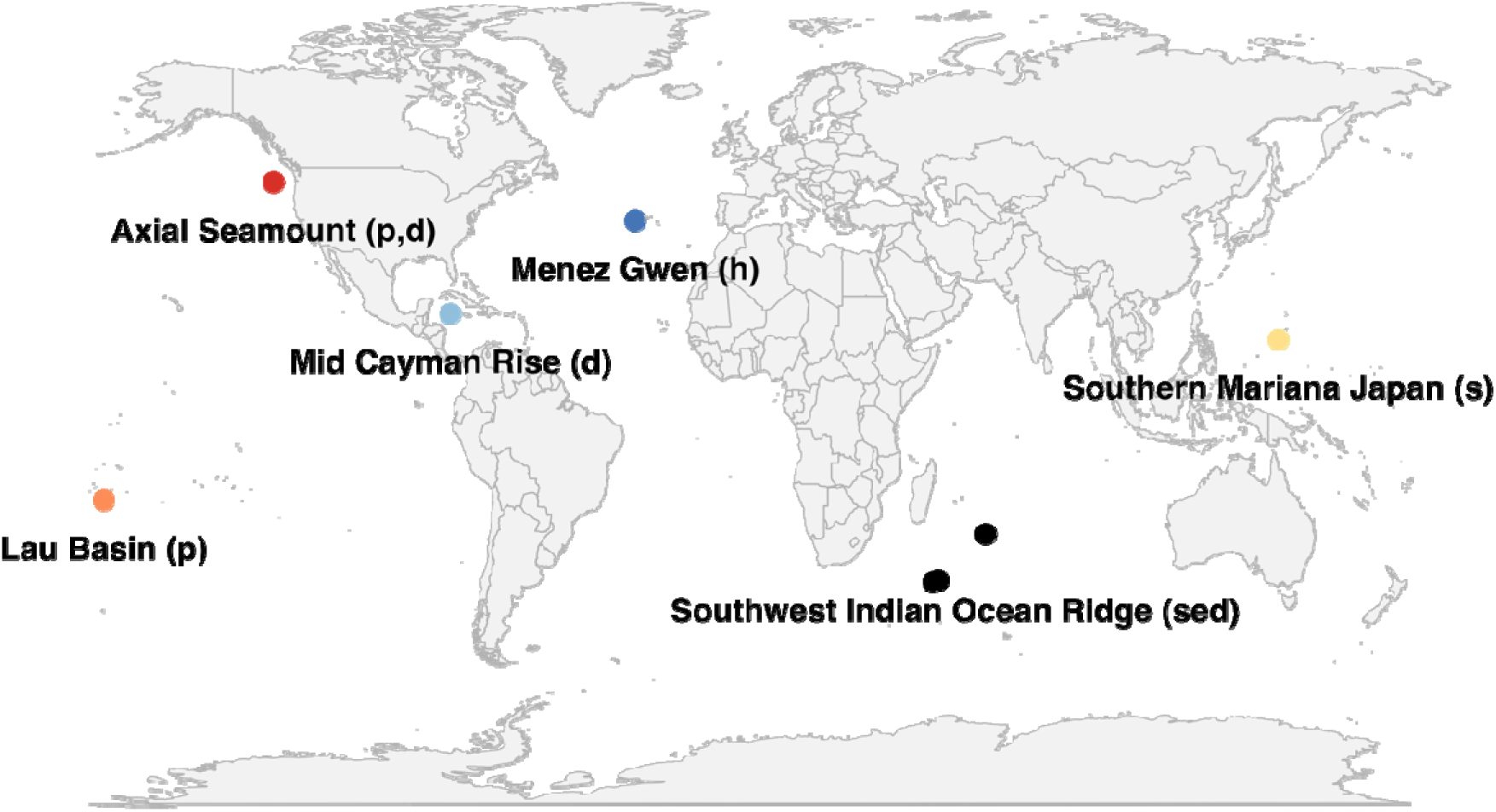
World map showing the global-distribution of metagenomic sampling locations analyzed in this study. Parentheses indicate the type of vent environment: d (diffuse fluid), h (hydrothermal fluid), p (plume), sed (sediment), and s (sulfide deposit).

**Fig. 2.**
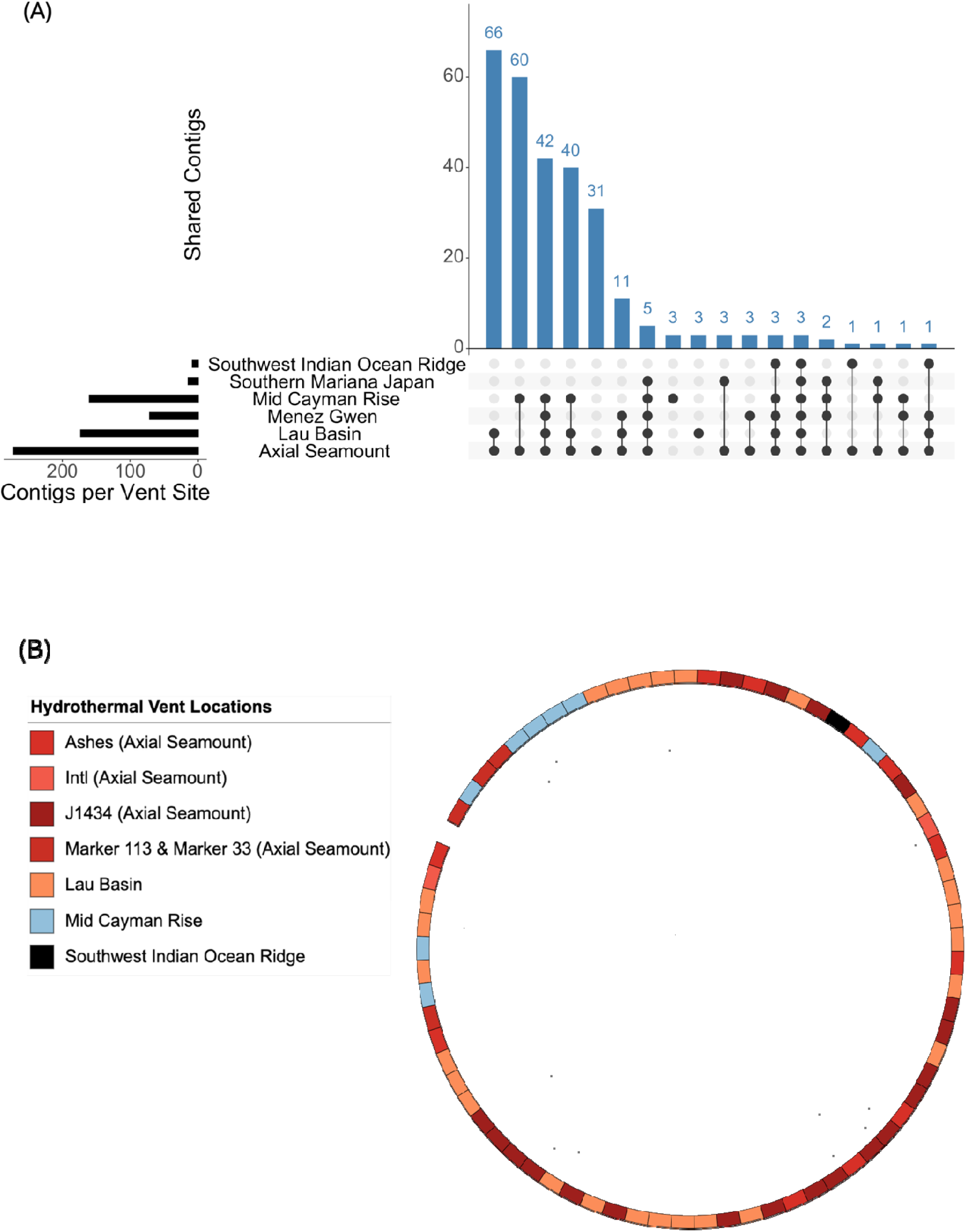
Distribution of NCLDV contigs across hydrothermal vents and their phylogenetic analysis. (A) Upset plot showing the number of shared NCLDV contigs detected across six hydrothermal vent locations. Black dots indicate the presence in one or multiple vent sites. Horizontal bar (left) shows the total number of NCLDV contigs detected in each vent site. The corresponding vertical bar shows the number of NCLDV contigs present in locations described by the black dots. (B) Phylogenetic tree (IQ-TREE model: LG+F+R5) of 73 NCLDV contigs based on a concatenated alignment of four NCLDV single copy marker genes (A32, PolB, VLTF3, SFII), visualized on iTOL. The outer ring indicates the hydrothermal vent location from which each contig was assembled.

t-SNE ordination revealed overlapping giant virus assemblages across samples in Axial Seamount (North Pacific), Lau Basin (South Pacific), and Menez Gwen (North Atlantic), suggesting similar giant virus community composition across these ocean basins (Fig. 3). Mid Cayman Rise samples formed distinct clusters, indicating a more unique community composition specific to this vent environment. Nevertheless, a subset of Mid Cayman Rise (Atlantic) samples overlapped with those from the Southern Mariana Japan (Equatorial Pacific), Southwest Indian Ocean Ridge, and Axial Seamount (North Pacific). Overall, the t-SNE ordination analysis revealed compositional overlap among hydrothermal vent giant virus communities, as well as regional distinctiveness across these major ocean basins.

**Fig. 3.**
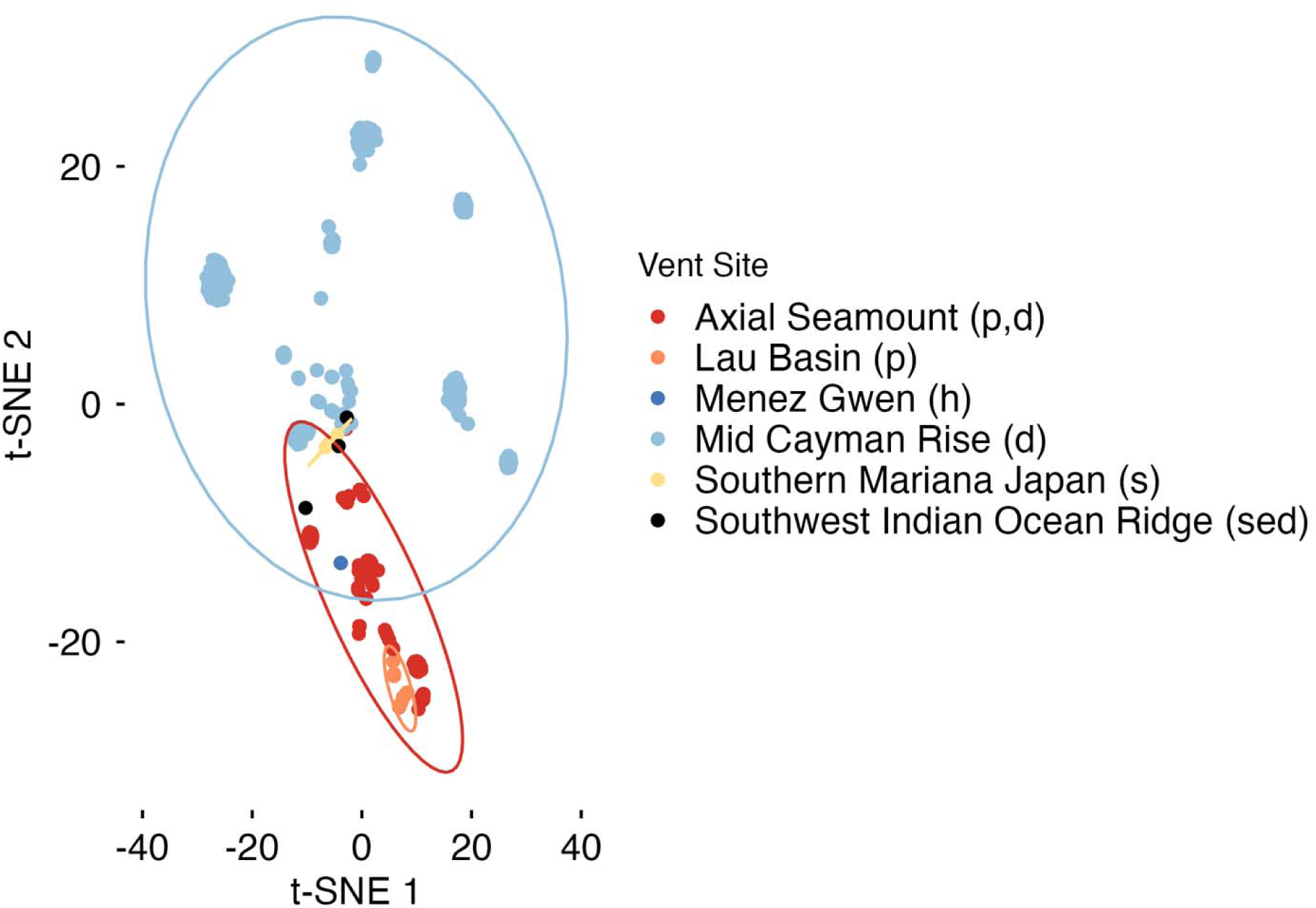
NCLDV community composition across geographically-distinct hydrothermal vents. t-SNE (t-Distributed Stochastic Neighbor Embedding) ordination plot based on the normalized Q2Q3 coverages of 279 NCLDV contigs across reads from 6 geographically-distinct vent locations. The interquartile coverage of each NCLDV contig was normalized to the lowest library size of metagenomic samples. Each dot represents one metagenomic sample and the color of the dot indicates the vent location. Parentheses indicate the type of vent environment: d (diffuse fluid), h (hydrothermal fluid), p (plume), sed (sediment), and s (sulfide deposit).

Our results corroborate previous reports suggesting the cosmopolitan nature and global distribution of giant viruses. For example, some lineages of giant viruses have been reported in metagenomic samples across terrestrial and marine environments [15]. In aquatic environments, the same giant virus OTUs were detected across distant and distinct aquatic ecosystems in Japan, with some of these same populations detected across multiple Tara Oceans samples obtained from the pelagic water column [19]. Biogeographic analysis of giant viruses across 480 samples obtained across the Atlantic and Pacific Oceans showed that several giant virus populations were particularly widespread and abundant in the oligotrophic pelagic water column [16]. Moreover, 101 deep-sea specific giant virus populations were identified on a global scale across 1890 metagenomic samples from 200 m up to 5601 m [20].

Our findings contrast with those from previous studies on hydrothermal vent viruses, primarily focused on bacteriophages, showing that viral populations are largely endemic to individual vent sites. For instance, a viral diversity study from Axial Seamount and Mid Cayman Rise reported that viruses are predominantly site-specific and not broadly distributed across vent systems [11]. Similarly, metagenomic analysis from eight vent sites across the global ocean found that viruses are highly endemic to vent sites and habitat types [12] . While vent-derived bacteriophage populations were mostly reported to be endemic, some viruses were found to be shared between geographically-distinct hydrothermal vent locations in the Pacific Ocean [10]. Taken together, these previous studies show that vent-derived bacteriophages are generally endemic and restricted to individual sites or a single ocean basin, whereas our results show that giant viruses can be broadly distributed across vent systems across distant ocean basins.

To investigate the evolutionary relatedness of the vent-derived giant virus populations across the global ocean, we performed a targeted phylogenetic analysis using four NCLDV single-copy marker genes [21]. Phylogenetic analysis using a concatenated tree showed that giant viruses from different hydrothermal vents clustered together, suggesting shared evolutionary lineages across globally-distributed vent systems (Fig. 2b). Phylogenetic clustering was most pronounced between contigs recovered from Axial Seamount (North Pacific) and Lau Basin (South Pacific), consistent with the observation that these locations shared the highest number of giant virus contigs (Fig. 2a). We also observed phylogenetic clustering between contigs recovered from these Pacific samples and Mid Cayman Rise (Atlantic), indicating that giant viruses can share evolutionary lineages across distant ocean basins. We then performed a comprehensive phylogenetic analysis of all 279 NCLDV contigs from six vent sites, using ten NCLDV-specific marker genes, which included multi-copy marker genes, to show that this pattern was consistent regardless of marker gene selection (fig. S5). Our observations are consistent with a previous study, which reported that giant virus OTUs from five geographically distant (between 74 and 1,765 km) and distinct aquatic ecosystems (coastal and offshore seawater, brackish water, and hot spring freshwater) in Japan also clustered together on the phylogenetic tree [19]. Our results show that giant viruses from the same evolutionary lineages or exhibiting close evolutionary relationships can occur in geographically distant hydrothermal vents, suggesting a shared evolutionary history amongst vent-derived giant virus populations across the globe.

Collectively, our biogeographic analysis showed a previously unexplored, wide distribution pattern of giant viruses across hydrothermal vents, distinct from the site-specific endemism that was previously observed for bacteriophages. We postulate that the broad distribution of phylogenetically similar giant viruses likely reflects 1. the presence of closely related giant virus lineages across vent systems, and/or 2. the broad functional capabilities of giant viruses, relative to the smaller genomes of bacteriophages, enabling the persistence of giant virus populations across ocean basins (“everything is everywhere”).

### Functional biogeography of vent-derived giant viruses

We postulate that the broad functional capabilities of giant viruses may contribute to their widespread distribution across deep-sea hydrothermal vents. Giant viruses contain large genomes, extending up to 2.5 million base pairs [22], that can encode an extensive repertoire of core and auxiliary functions [23]. To study the functional biogeography of giant viruses, we examined the biogeographic patterns of predicted protein functional domains from PFAM across six vent systems. A total of 181 PFAM domains were unique to the Axial Seamount, consistent with expectations from a higher number of endemic giant virus populations in this location (fig. S6, Fig. 2a). The highest number of PFAM domains was shared among Axial Seamount (North Pacific), Lau Basin (South Pacific), and Menez Gwen (North Atlantic). Eleven PFAM domains were present in all vent locations, all of which are associated with DNA-dependent RNA polymerase (fig. S6). RNA polymerase is encoded by nearly all giant viruses [23,24] and enables them to complete the infection cycle within the host cytoplasm [25]. 59 PFAM domains were detected ≥4 out of the six vent systems (table S4), including ribonucleotide reductase and tRNA synthetase. Ribonucleotide reductases are widely present among members of the *Nucleocytoviricota* [26] and catalyze the conversion of ribonucleotides to deoxyribonucleotides, a key step in DNA synthesis and repair [27]. Similarly, giant viruses encode components of protein translation machinery, including aminoacyl tRNA synthetase [28]. The broad biogeographic distribution of these giant virus–encoded functional domains, which support fundamental processes associated with replication, DNA synthesis, and translation, reveals the core functional machinery required to inhabit diverse hydrothermal vent environments.

To look for evidence of local adaptation, we identified a key gene that may facilitate adaptation to specific hydrothermal vent habitats. Heat-shock proteins (HSPs) were previously identified in giant viruses in both culture-based and metagenomic studies [16,29,30]. In hydrothermal vents, several eukaryotic species were found to express HSPs to cope with the heat stress [31,32]. We identified giant viruses encoding HSPs in three hydrothermal vent systems, with particularly high relative abundances in Axial Seamount samples (fig. S7). Among individual sites within Axial Seamount, the highest relative abundance of HSP-containing giant viruses was observed in J1434, a diffuse fluid sample that originated from the subsurface, characterized by elevated temperature (∼50°C) compared to other samples from this location. The enrichment of HSP-encoding giant viruses in high-temperature diffuse fluids may reflect local thermal adaptation to hot diffuse fluids originating from the subsurface. This result is consistent with local adaptation (“the environment selects”) and the idea that increased functional versatility, due to larger genomes, may contribute to the wide biogeographic distribution of giant viruses across hydrothermal vent ecosystems.

### Passive transport via hydrothermal plumes drives the biogeography of giant viruses across the global ocean

To investigate the distribution of hydrothermal vent–derived giant viruses in the global ocean, we further analyzed 868 publicly available pelagic metagenomic samples collected from 5 m to 4000 m across geographically-distinct pelagic samples [33–38] (table S5). On average, 0.37% of reads from global ocean metagenomes mapped to the 279 NCLDV contigs recovered from hydrothermal vents (Fig. 4). Endemic giant viruses, defined as populations detected at only a single vent site, showed the lowest relative abundances across pelagic ocean samples. In contrast, widely distributed giant viruses, defined as presence across four or more of the six vent sites, showed the highest relative abundances across pelagic ocean samples (fig. S8). This pattern is consistent with the idea that globally-distributed giant virus populations, which are observed across >=4 or more hydrothermal geographically-distinct vents systems, are transported via the pelagic ocean. Although the percentage of reads mapped to vent-derived NCLDV contigs across pelagic samples are low (0.37%), they are similar compared to percent reads mapped to NCLDV contigs obtained from the same environment, which range from to 0.03–1.5% in vent samples analyzed here and ∼0.5% in pelagic open ocean samples [26]. This comparison suggests that our results were not a byproduct of false-positives in read-mapping, and instead indicate that vent-derived giant viruses were indeed present in pelagic ocean samples. Observing vent-derived giant viruses across the global pelagic ocean could also support the microbial seed bank hypothesis [39], with giant viruses in the pelagic ocean potentially fueling their colonization across hydrothermal vents. Both of these processes, horizontal passive transport and the seed bank hypothesis, could occur simultaneously to result in the global distribution and local selection of giant virus populations.

**Fig. 4.**
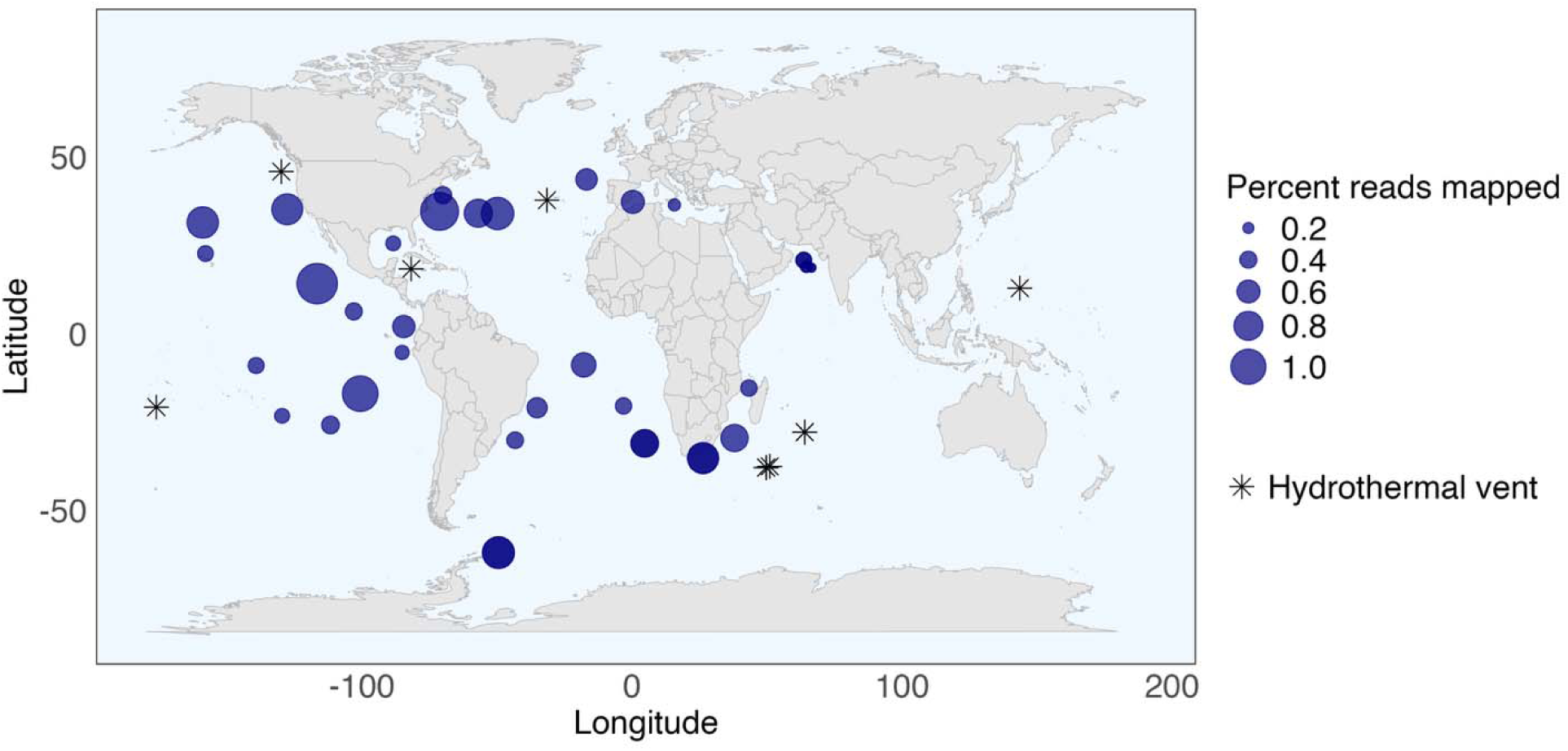
Distribution of vent-derived giant viruses in global ocean. Percentage of reads mapped (navy circles) to 279 non-redundant NCLDV contigs from hydrothermal vents across global ocean pelagic metagenomes at depths ranging from 5 to 4000 m, averaged across depths for each location. The asterisks indicate hydrothermal vent locations where NCLDV contigs were assembled.

The highest percentages of pelagic reads mapping to vent–derived giant viruses were observed in the North and South Pacific Oceans and the North Atlantic Ocean (Fig. 4, table S6).

Furthermore, a ∼3-fold higher percentage of pelagic reads from mesopelagic samples (350–4000 m), compared to upper ocean samples, mapped to hydrothermal plume-derived giant viruses (Fig. 5). This observation is consistent with expectations from stratification in the upper ocean, which prevents vertical mixing, and the horizontal passive transport of vent–derived giant viruses via vent plumes across the mesopelagic ocean. Taken together, these observations suggest that horizontal passive transport, rather than the seed bank hypothesis, may play a more significant role in the distribution of giant viruses across our global ocean.

**Fig. 5.**
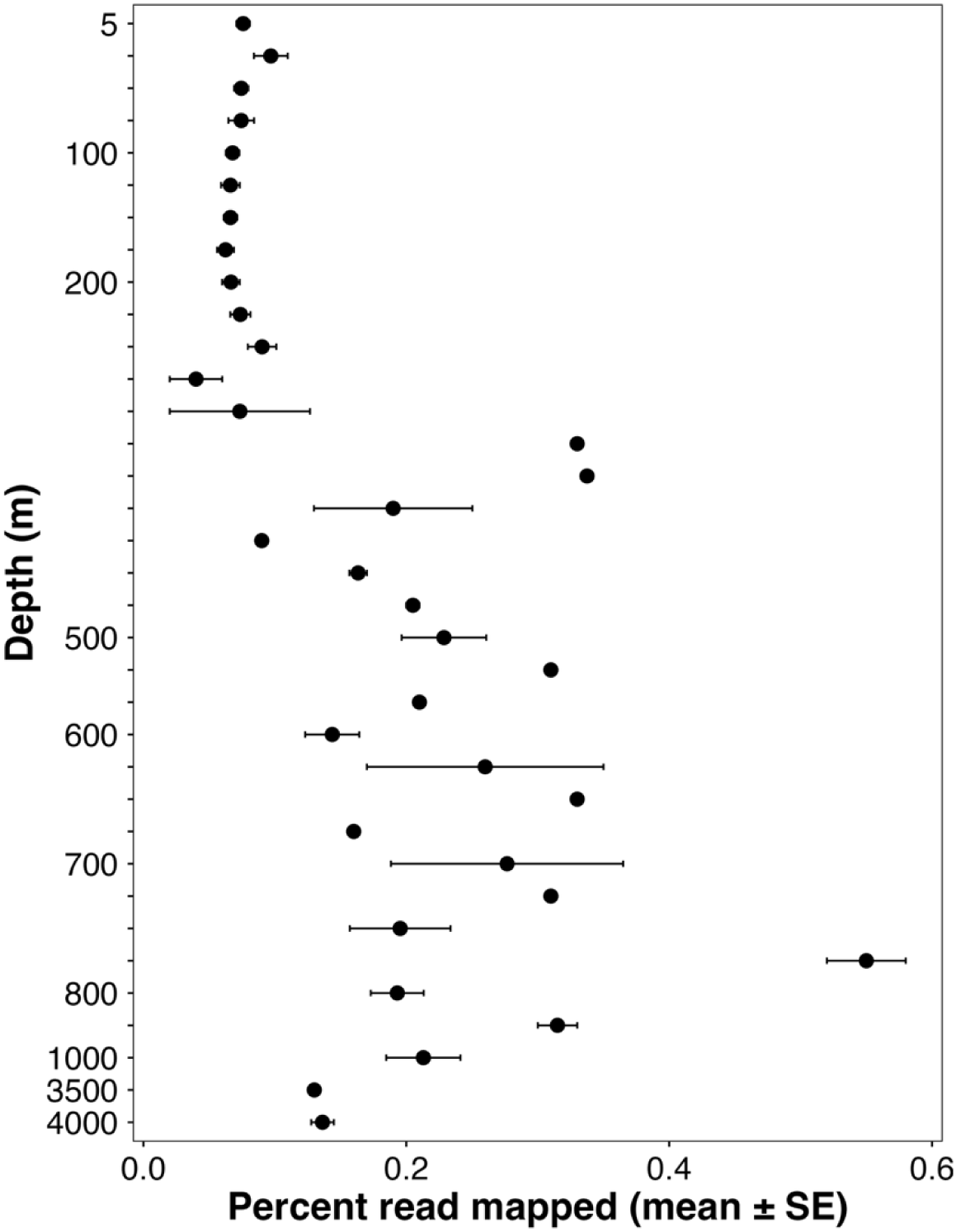
**Depth distribution (percent reads mapped) of plume-derived NCLDV contigs across global ocean metagenomes**. Percent reads mapped to NCLDV contigs are averaged across all locations for each depth. Plume-derived NCLDV contigs (n=133), defined as those assembled from hydrothermal plume samples (Lau Basin, Ashes and Intl from Axial Seamount), were extracted from a total 279 nonredundant NCLDV contigs for read-mapping.

Hydrothermal vent plumes may act as underwater highways, facilitating the long-distance dispersal of microbial communities across hundreds of kilometers [40]. Hydrothermal plumes rise upon discharge into seawater due to their lower density and progressively dilute as they ascend, eventually reaching neutral buoyancy [5]. Once neutral buoyancy is achieved, the plume water masses disperse laterally through the water column, following ocean currents [5].

Microbial communities associated with vent plumes have been detected well beyond vent fields, including in mesopelagic and bathypelagic habitats [41], suggesting that hydrothermal plumes may help establish ecological connections between pelagic and seafloor/subsurface habitats.

Vent-derived bacteria were also present in the open ocean water globally and they can potentially be transported over large distances across the water column through vent plumes [2].

Furthermore, planktonic larvae can be dispersed and transported long distances through deep ocean currents [42]. The South Equatorial Current has been proposed as a mechanism driving larval dispersal across the southwest Pacific vent complex spanning more than 4,000 km [43], and gastropod larvae can travel over 350 km before settling at vents along the East Pacific Rise [44]. Couched in this broader context, our study indicates that vent–derived giant viruses are not confined to their source habitats and may be broadly distributed across the globe through horizontal passive transport in the mesopelagic ocean.

## Conclusion

We found that the hypothesis “everything is everywhere, but the environment selects” holds true for giant virus populations inhabiting deep-sea hydrothermal vents. The biogeography of giant viruses showed wide distribution across hydrothermal vents from distinct ocean basins, contrasting with the site-specific endemism previously observed in vent-derived bacteriophages. We observed evidence suggesting that this widespread distribution reflects shared evolutionary lineages across geographically-distant vent systems and the functional versatility encoded by their large genomes. We show that this global distribution is likely driven by horizontal passive transport via hydrothermal plumes. Our study also indicates that environmental selection can shape the distribution of giant viruses within vent ecosystems, shown by evidence supporting local thermal adaptation. Taken together, these results indicate that the global distribution of giant virus populations are driven by horizontal passive transport via hydrothermal plumes. Our study shows that a classic paradigm in microbial biogeography applies to marine giant viruses, and identifies the mechanisms maintaining the dispersal and spatial ecology of giant viruses across the global ocean.

## Methods

### Sample collection, DNA extraction and metagenomic sequencing

Diffuse vent fluid and neutrally buoyant plume samples were collected from Axial Seamount (45°58’ N 130°00’ W), located on the Juan de Fuca Ridge 480km west of the Oregon coast (USA), during research cruise R/V Thomson in July 2022. Plume samples were collected ∼50–250m above the seafloor at the Axial Seamount Hydrothermal Emission Study (ASHES) and International District (Intl) using Niskin bottles connected to a CTD, while diffuse fluid was sampled at ASHES-Anemone site using the ROV Jason (J1434). A total of 2300 mL from ASHES, 1060 mL from International District, and 500 mL from J1434, respectively per sample, was filtered through a 0.22 µm Sterivex filter using a peristaltic pump. Microbial DNA was extracted from the filter using Masterpure 659 extraction kit (Lucigen MC85200), followed by library preparation using the Illumina KAPA 687 HyperPrep kit and sequencing on the Illumina Novaseq X plus 10B platform.

### Metagenomic datasets

To investigate the biogeography of giant viruses in deep-sea hydrothermal vents, we obtained and analyzed an additional 511 publicly available metagenomic samples collected from six globally-distributed vent systems. These samples included 145 diffuse fluid samples from Axial Seamount in the Pacific Ocean [45], 343 diffuse fluid samples from the Mid Cayman Rise in the Atlantic Ocean [11], 11 plume samples from the Lau Basin in the Pacific Ocean [46], 3 hydrothermal fluid samples from Menez Gwen [47], 4 sulfide deposit samples from the Southern Mariana (Japan) region in the Pacific Ocean [48], and 5 sediment samples from the Southwest Indian Ridge [49]. All samples were derived from the >0.2 µm cellular size fraction, as giant viruses are expected to be present within this range. Geographic locations and metadata for these hydrothermal vent datasets are available on Fig. 1 and table S1, respectively. Technical details of the bioinformatic workflow can be found in fig. S1.

### Data preprocessing and assembly

The quality of the raw reads was assessed through FastQC v.0.11.9 [50] and MultiQC v.1.21 [51]. Raw reads were further trimmed using Trimmomatic v.0.39 [52] and BBDuk v.39.01[53] to remove the adapter sequence and low-quality bases. Next, cleaned reads were assembled into contigs using the de novo assembler MegaHit v.1.2.9 [54] using a minimum contig length of 5000 bp. Cleaned reads from Ashes (Axial seamount), International District (Axial seamount), J1434 (Axial seamount), Lau Basin, Southern Mariana Japan, and Southwest Indian Ocean Ridge were coassembled. Cleaned reads from publicly available Axial Seamount and Mid Cayman Ridge samples were assembled separately for each metagenome due to the large number of samples. An initial NCLDV-specific reassembly was performed for publicly available Axial Seamount and Mid-Cayman Ridge metagenomic samples. Briefly, all the generated contigs from publicly available Axial Seamount and Mid Cayman Ridge were subsequently analyzed through Virsorter2 v2.2.4 [55] using the parameter --include-groups "dsDNAphage,NCLDV,RNA,ssDNA,lavidaviridae". Contigs classified as NCLDV were extracted and reads mapping to these contigs were identified using Bowtie2 v.2.5.1[56] and SAMtools v.1.19 [57]. Finally, all mapped reads to Virsorter2 classified NCLDV contigs were pooled to perform NCLDV-specific reassembly using MegaHit v.1.2.9 [54] for publicly available Axial Seamount and Mid Cayman Ridge samples. In total, 308,894 contigs (>5kbp) were assembled from all hydrothermal vent locations.

### Generation of non-redundant NCLDV contig database

All 308,894 assembled contigs were analyzed through ViralRecall v.2.0 [58], which identifies NCLDV contigs based on the homology to known NCLDV proteins. We identified 442 contigs (>5kbp) containing at least one of ten NCLDV-specific marker genes (E-value ≤ 1e-05) and a ViralRecall score ≥ 3 [59]. A positive ViralRecall score indicates a viral origin, whereas negative scores suggest possible cellular contamination. The ten NCLDVs marker genes used were NCLDV major capsid protein (MCP), superfamily II helicase (SFII), virus-like transcription factor (VLTF3), B-family DNA polymerase (PolB), and A32-like ATPase (A32), DNA-dependent RNA polymerase alpha subunit (RNAPL), DNA-dependent RNA polymerase BETA subunit (RNAPS), transcription elongation factor II-S (TFIIS), DNA helicase primase (D5), and ribonucleotide reductase (RNR). The NCLDV contigs were dereplicated using cd-hit-est v.4.8.1

[60] at >95% ANI across >90% of the shorter contig. These steps resulted in 279 nonredundant NCLDV contigs longer than 5 kbp, containing a total of 521 NCLDV-specific marker genes, with a mean ViralRecall score of 6.88, indicating a strong viral origin.

### Taxonomic and functional annotation of NCLDV contigs

The taxonomic identification of NCLDV contigs was performed using the NCLDV marker proteins reported above. Briefly, the marker proteins identified in the NCLDV contigs were subjected to BLASTp v.2.11.0+ [61] against the protein sequences from the giant virus database [62] with an E-value of 1 × 10-5, a percent identity of >60% and a query coverage of >50%.

Functional annotation of the NCLDV contigs was performed using the Pfam database (release 36.0) and HMMER v3.3.2 [63], applying the --cut_nc parameter to assign protein domains based on curated PFAM noise cutoffs.

### Phylogenetic analysis of NCLDV contigs

Phylogenetic analysis of the NCLDV contigs based on four NCLDV single-copy marker genes, including A32, PolB, VLTF3, and SFII [21], was performed to investigate the evolutionary relationships of contigs originating from geographically distinct hydrothermal vent locations.

Briefly, proteins encoded by NCLDV contigs were predicted using Prodigal v.2.6.3 [64]. Multiple sequence alignment for each of the four NCLDV single-copy marker genes was generated using mafft v7.525woe [65], followed by concatenating alignment with AMAS v.1.0 [66]. The alignment was further trimmed with trimAI v.1.4.1 [67] (parameter -gt 0.1). IQ-TREE v2.3.0 [68] was utilized to develop a maximum-likelihood phylogenetic tree with 1000 ultrafast bootstraps (model: LG+F+R5). The tree was visualized by the interactive tree of life (iTOL) v.7.4 [69]. An additional phylogenetic tree using all ten previously mentioned NCLDV-specific marker genes was constructed following the same approach with 1000 ultrafast bootstraps (model: LG+R7).

### Geographic distribution of NCLDV contigs in hydrothermal vents

To investigate the distribution of NCLDV contigs across hydrothermal vents, quality-controlled reads from all the hydrothermal vent metagenomic samples were mapped on 279 nonredundant NCLDV contigs (>5kbp) using BWA v.0.7.17 [70] with default parameters. Next, the interquartile mean coverage of NCLDV contigs was calculated for each sample using Anvi’o v.8 [71]. The interquartile coverage provides more accurate estimates of coverage because it reduces the overestimation of coverage in conserved regions and the underestimation in hypervariable regions. For each contig, its interquartile coverage was normalized to the lowest library size of metagenomic samples, and contigs with coverage above zero were considered to be present in the sample. t-SNE (t-Distributed Stochastic Neighbor Embedding) was performed using the normalized interquartile coverages of NCLDV contigs to visualize variability in the composition of NCLDV assemblages across hydrothermal vents. The relative abundance of NCLDV contigs was calculated by normalizing the interquartile of a contig by the total interquartile coverage in that sample.

### Distribution of hydrothermal vent NCLDV contigs in the global pelagic ocean

To investigate the global distribution of hydrothermal vent NCLDV contigs across pelagic samples, we analyzed 868 previously sequenced metagenomic samples [33–38] spanning depths from 5m to 4000 m. These samples included mesopelagic Tara Oceans samples (270–1,000m, n = 68), Station ALOHA samples (5–4,000m, n = 798), Mediterranean Sea samples (1,000 and 3,000 m, n = 2). Reads from 868 metagenomic samples were mapped to nonredundant NCLDV contigs from hydrothermal vents using BWA v.0.7.17 [70] with default parameters. The percentage of reads mapped from each sample to hydrothermal vent NCLDV contigs was calculated using SAMtools v.1.19 [57]. From a set of 279 nonredundant NCLDV contigs, 133 contigs assembled from hydrothermal plume samples (Lau Basin, Ashes, and International at Axial Seamount) were extracted and reads from 868 pelagic metagenomic samples were mapped to nonredundant contigs assembled from hydrothermal plume.

## Acknowledgments

We thank R/V Thompson, TN405, ROV Jason crew on dive J1434, Chief Scientist Julie Huber, Sarah Hu, Maria Pachiadaki, (cruise funded by National Science Foundation: OCE-1947776 to JH, MP, SH). Sabrina Elkssas, Bayleigh Benner. We thank Paulo Freire for performing quality checks and providing assemblies of the collected Axial Seamount reads. We also acknowledge Rebekah Rogers, Elizabeth Cooper, Abigail Labella, Way Sung for their discussion on phylogenetic analysis.

## Author contributions

EL conceptualized, supervised, and acquired funding for this study. MMS performed the analysis. MMS and EL wrote the manuscript.

## Data and materials availability

The raw metagenomic sequence reads of three collected Axial Seamount and NCLDV assemblies will be available publicly in the National Center for Biotechnology Information (NCBI) upon publication .

---

**Fig. S1.**
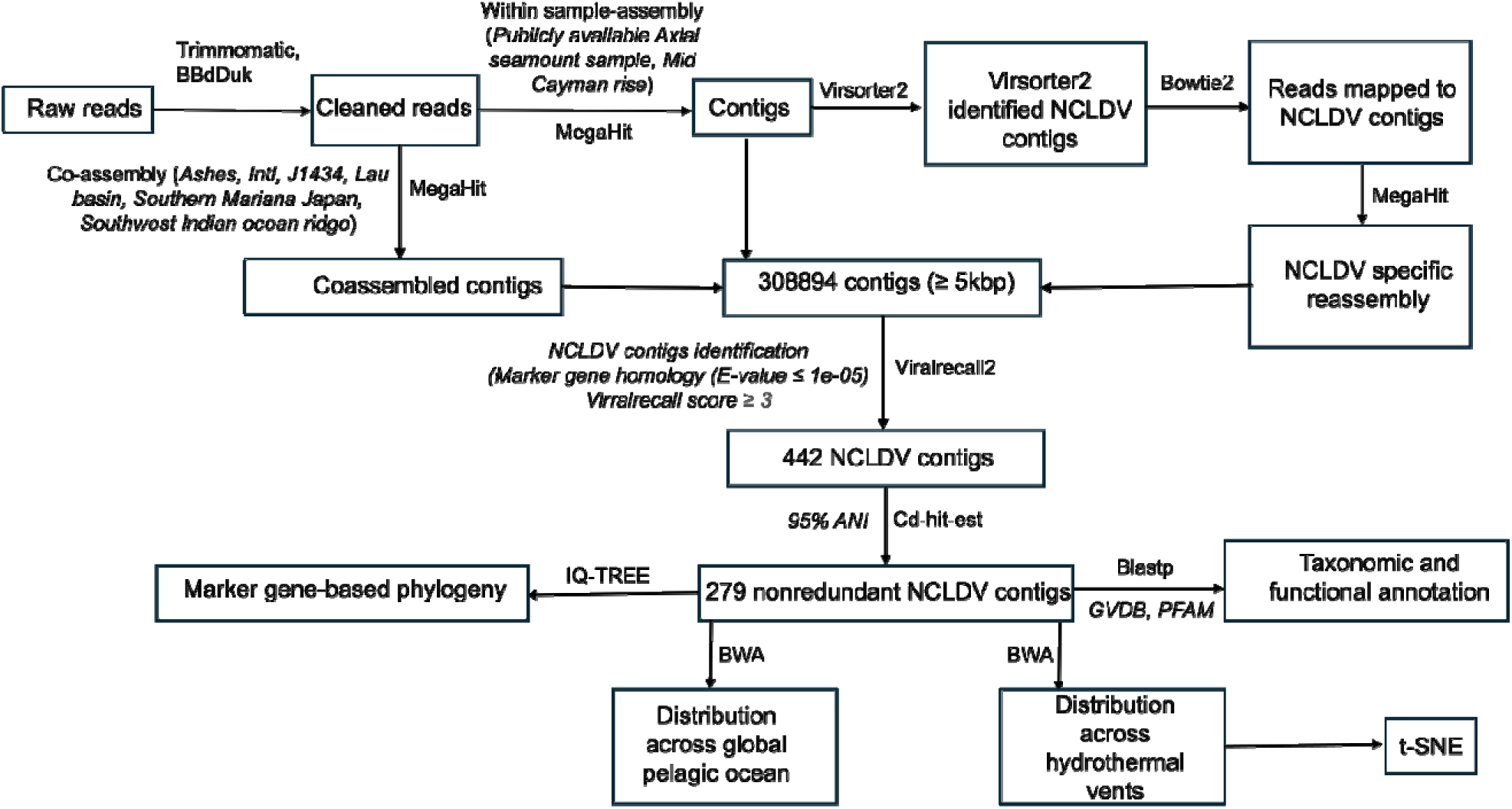
Bioinformatic pipeline utilized in this study. Schematic workflow from raw metagenomic reads to NCLDV contigs database and analysis from 514 metagenomic samples including three samples collected in this study and 511 samples previously collected across 6 geographically-distinct vents sites .

**Fig. S2.**
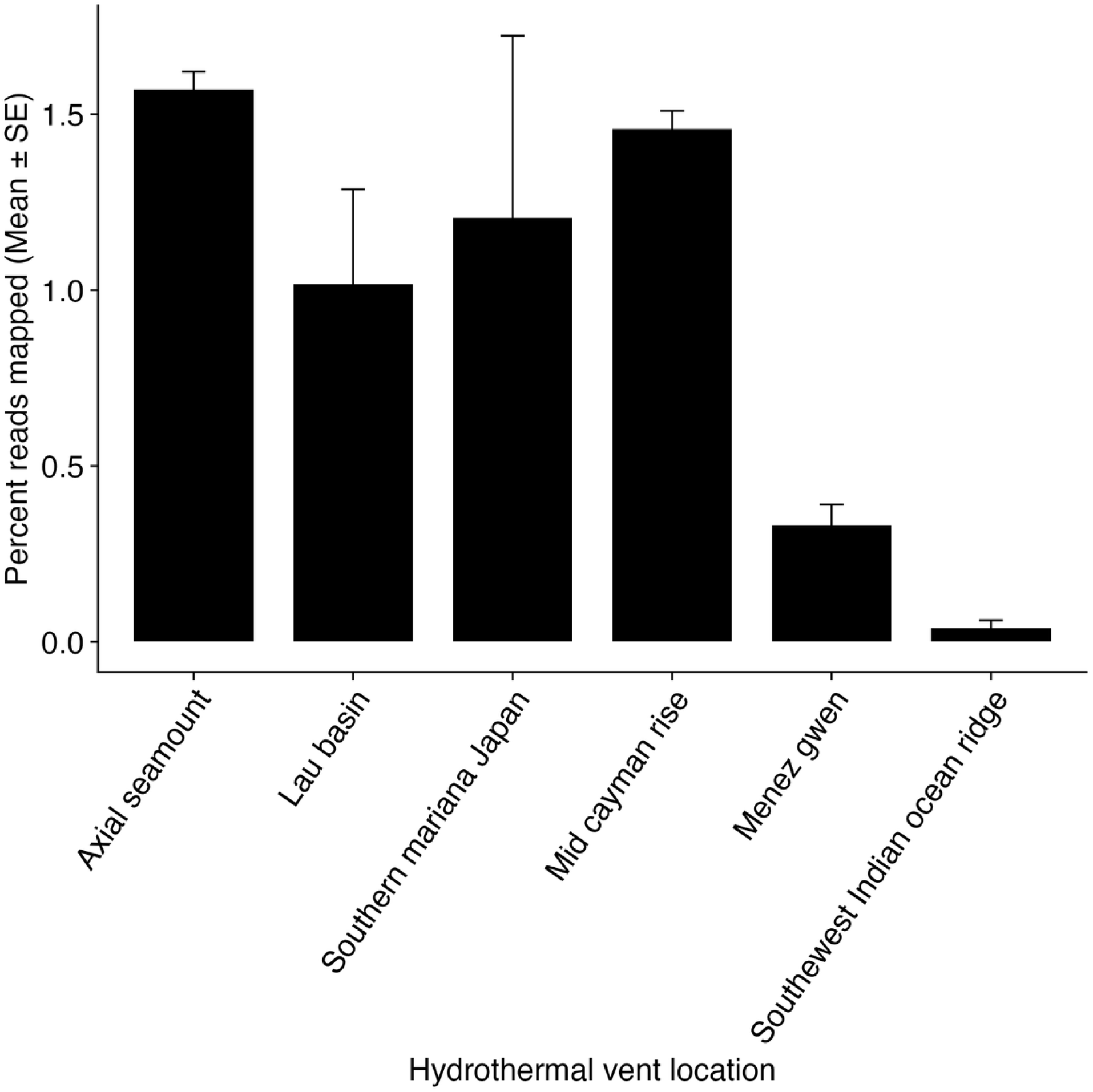
**Percentage of reads mapped (mean + SE) to 279 nonredundant NCLDV contigs across geographically-distinct hydrothermal vent locations**. The mean was calculated across each of the six vent sites, and error bars indicate standard error (SE).

**Fig. S3.**
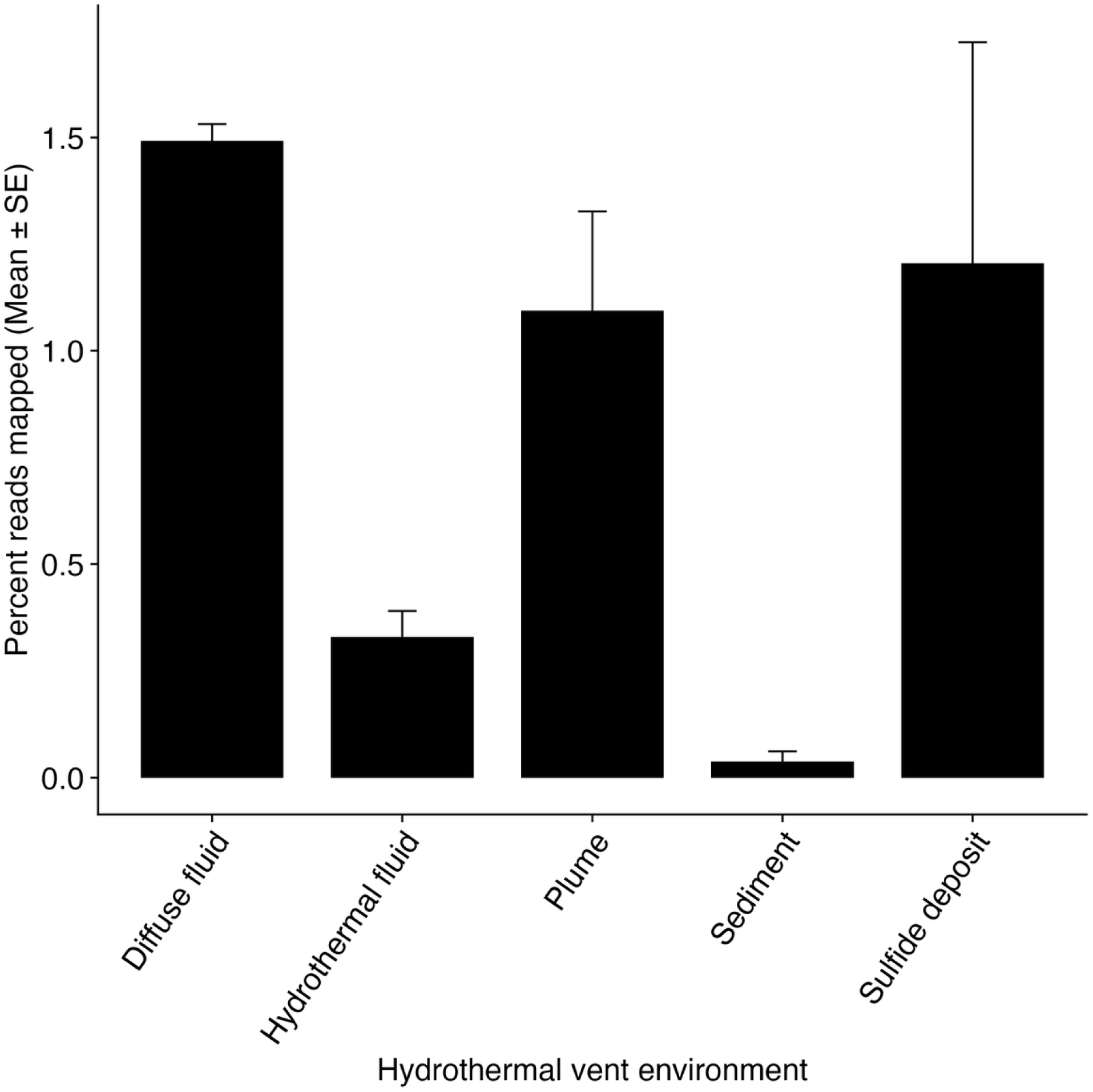
**Percentage of reads mapped (mean + SE) to 279 nonredundant NCLDV contigs across hydrothermal vent environments**. The mean was calculated across each of the five vent environments, and error bars indicate standard error (SE).

**Fig. S4.**
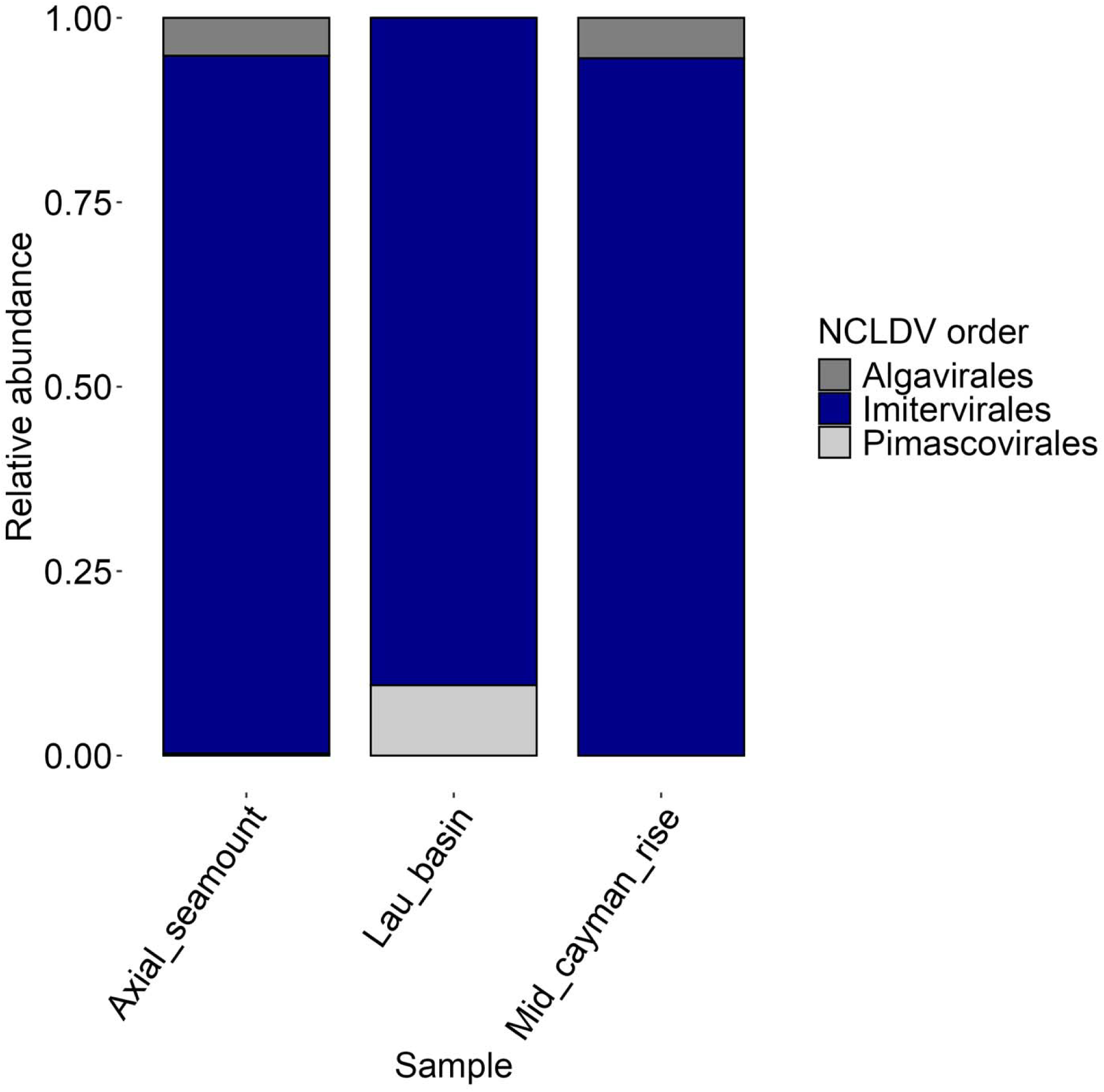
**Relative abundances of taxonomically identified NCLDV contigs, grouped by their taxonomic assignments**. Order level taxonomy was assigned through marker gene-based homology with the giant virus database. Relative abundances were calculated using interquartile coverages of contigs normalized to the maximum coverage of all contigs in that group for those samples. The legend shows the order-level taxonomic classification of NCLDV contigs

**Fig. S5.**
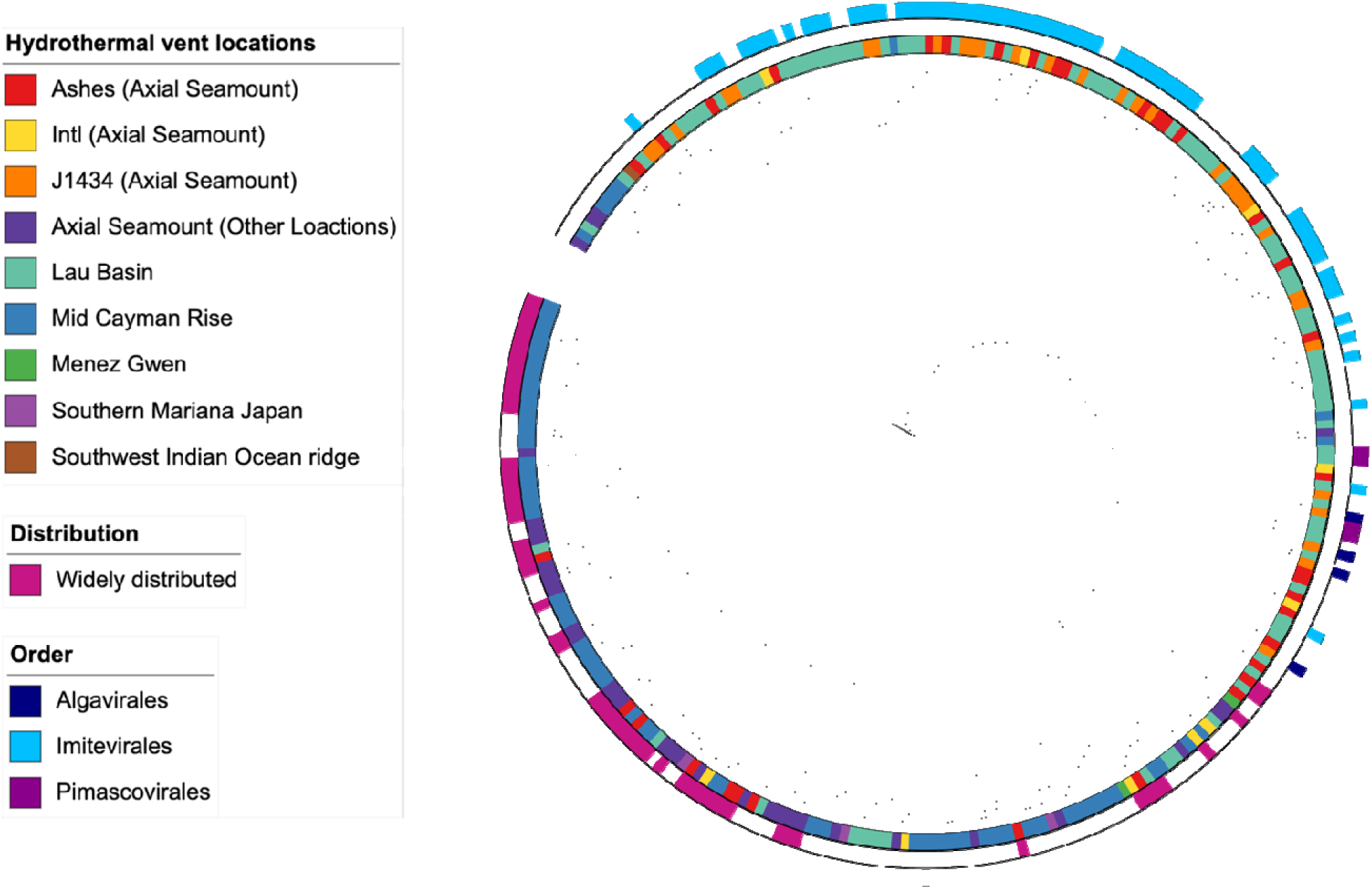
Phylogenetic analysis of the vent-derived NCLDV contigs. Phylogenetic tree (IQ-TREE model: LG+R7) of 279 NCLDV contigs based on a concatenated alignment of ten NCLDV marker genes (A32, PolB, VLTF3, SFII, mcp, RNR, RNAPS, RNAPL, D5, TFIIS), visualize on iTOL [69]. The inner ring indicates the hydrothermal vent location from which each contig was assembled. The middle ring indicates widely-distributed giant virus contigs, defined by their presence in ≥4 out of the six vent systems and the outer ring indicates the order-level taxonomy of the NCLDV contigs, as identified through marker gene based homology with the giant virus database [62].

**Fig. S6.**
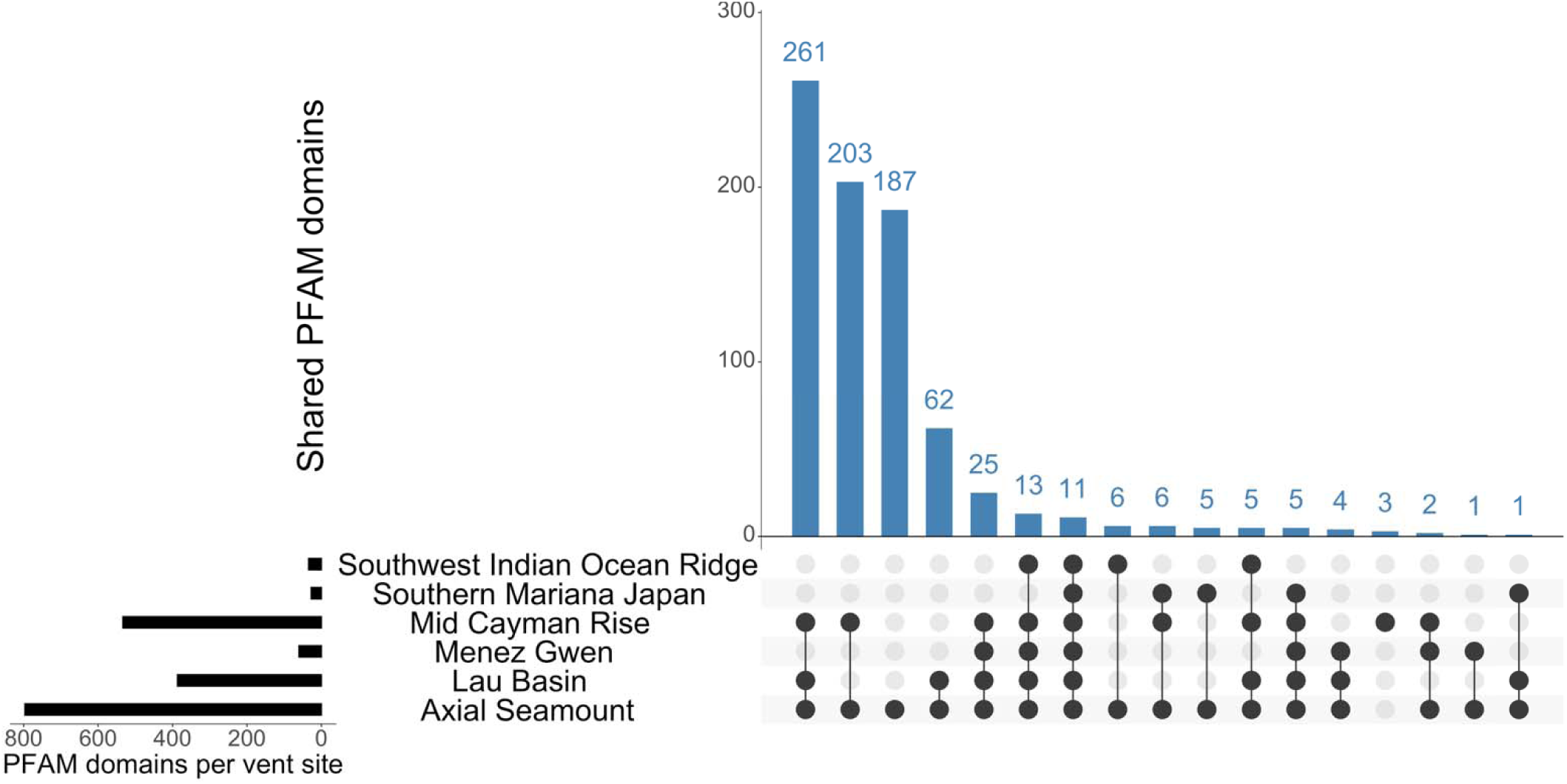
Number of shared PFAM domains detected across six hydrothermal vent locations. Black dots indicate the presence in one or multiple vent sites. The horizontal bar (left) shows the total number of PFAM domains detected in each vent site. The corresponding vertical bar shows the number of PFAM domains shared across locations connected by the black dots

**Fig. S7.**
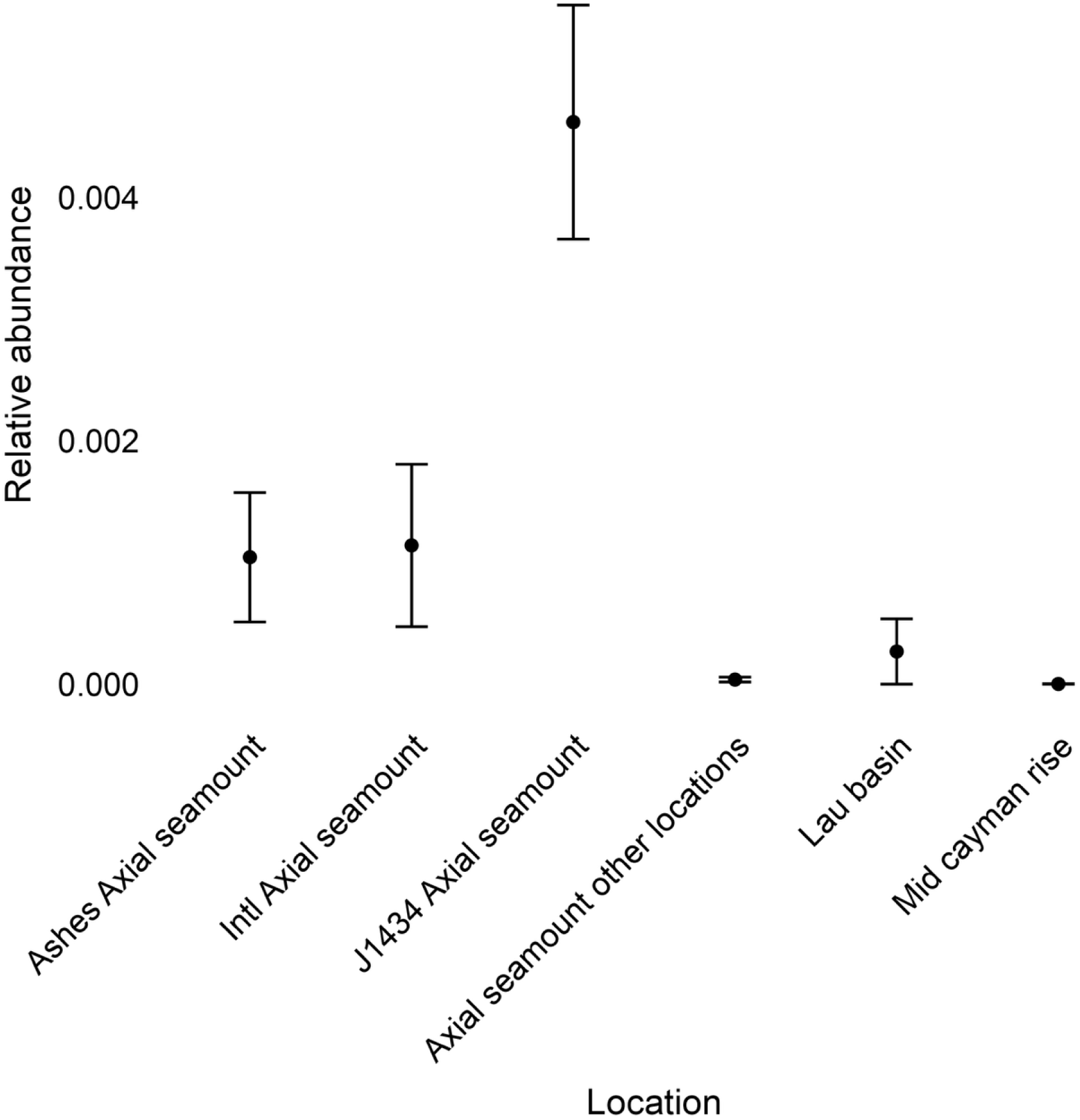
Relative abundances (mean +/- SE) of NCLDV contigs containing heat-shock proteins across vent sites. Relative abundance was calculated using interquartile coverage normalized to the total interquartile coverage of all NCLDV contigs in a sample.

**Fig. S8.**
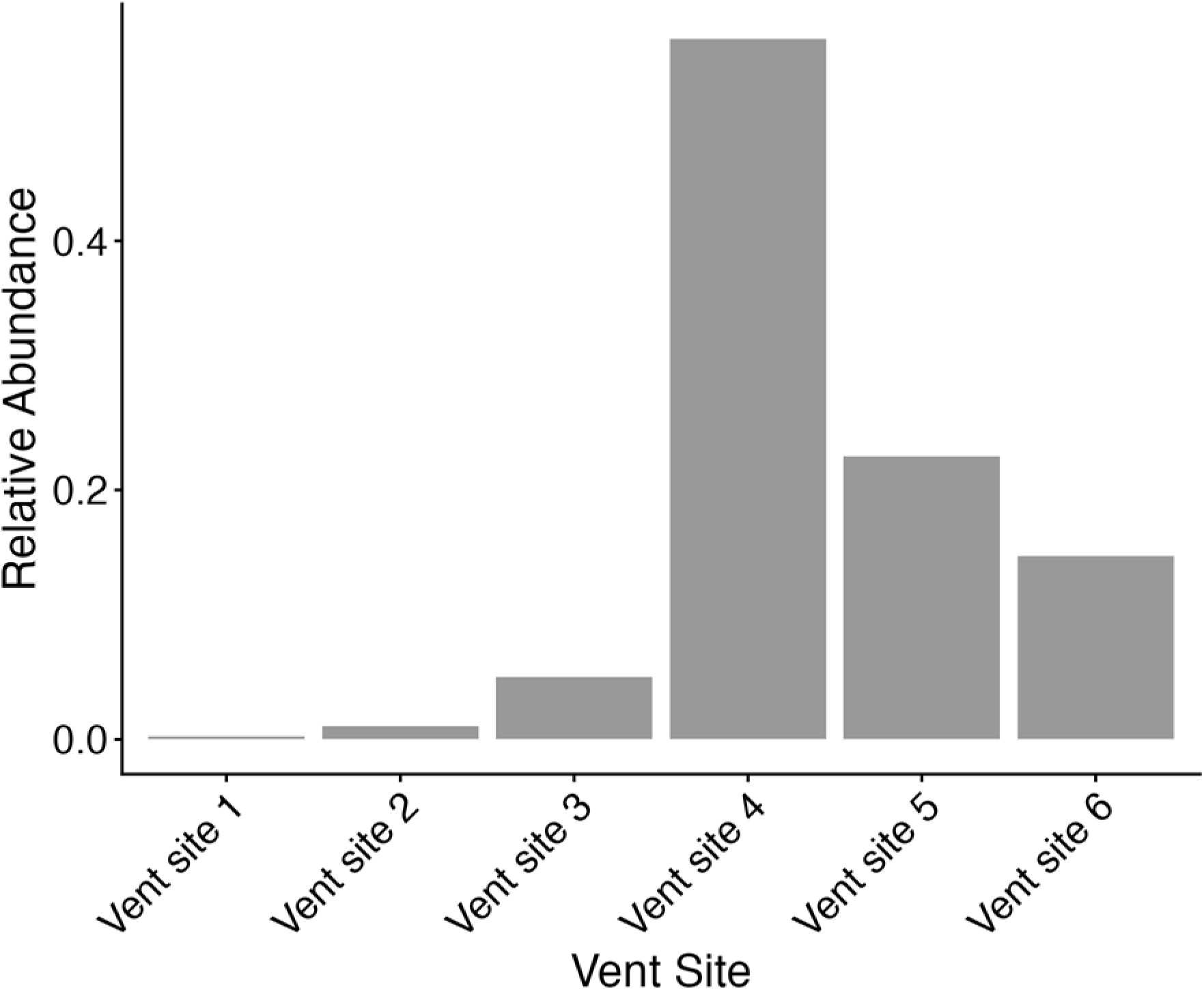
Relative abundance of endemic and widely distributed hydrothermal vent giant viruses across all pelagic samples from the global ocean. The x-axis shows the number of hydrothermal vent sites in which giant virus contigs were detected, with site 1 representing endemic contigs and site 6 representing contigs distributed across all vent sites. Relative abundances was calculated for every pelagic sample using interquartile coverage of each NCLDV contig, normalized to the total interquartile coverage of all NCLDV contigs in the pelagic sample

